# stMMR: accurate and robust spatial domain identification from spatially resolved transcriptomics with multi-modal feature representation

**DOI:** 10.1101/2024.02.22.581503

**Authors:** Daoliang Zhang, Na Yu, Wenrui Li, Xue Sun, Qi Zou, Xiangyu Li, Zhiping Liu, Zhiyuan Yuan, Wei Zhang, Rui Gao

## Abstract

Deciphering spatial domains using spatially resolved transcriptomics (SRT) is of great value for the characterizing and understanding of tissue architecture. However, the inherent heterogeneity and varying spatial resolutions present challenges in the joint analysis of multi-modal SRT data. We introduce a multi-modal geometric deep learning method, named stMMR, to effectively integrate gene expression, spatial location and histological information for accurate identifying spatial domains from SRT data. stMMR uses graph convolutional networks (GCN) and self-attention module for deep embedding of features within unimodal and incorporates similarity contrastive learning for integrating features across modalities. Comprehensive benchmark analysis on various types of spatial data shows superior performance of stMMR in multiple analyses, including spatial domain identification, pseudo-spatiotemporal analysis, and domain-specific gene discovery. In chicken heart development, stMMR reconstruct the spatiotemporal lineage structures indicating accurate developmental sequence. In breast cancer and lung cancer, stMMR clearly delineated the tumor microenvironment and identified marker genes associated with diagnosis and prognosis. Overall, stMMR is capable of effectively utilizing the multi-modal information of various SRT data to explore and characterize tissue architectures of homeostasis, development and tumor.

## Introduction

The advancement in spatially resolved transcriptomics (SRT) technologies has opened new avenues for a deeper understanding of the spatial architecture and functionality of tissues. By employing these advanced SRT techniques, we can conduct a precise examination of the spatial landscapes and transcriptional profiles for complex tissues. Currently, many SRT technologies have been developed, such as imaging-based and sequencing-based methods (1–7). Among these, techniques such as 10x Genomics Visium not only provide the spatial location and gene expression data for each spot but also acquire high-resolution hematoxylin and eosin (H&E) stained histology images of the tissue section, revealing richer information about the tissue organization. These technological advancements offer new insights into characterization of tissue architecture, enabling a more comprehensive understanding of tissue development and disease pathogenesis (7–9).

For SRT technologies capable of providing both gene expression data and histology images, the information from these different modalities reflects the structural information of tissues at various levels. Gene expression profiles reflect the difference of cell state between spots (10). Spatial location information provides the precise location of each spot, illustrating the complex landscape of tissue structure. Histological images display morphological features of cells, such as size, shape, and intercellular relationships, offering crucial clues for distinguishing functional domains (11). Although each of these modalities has its own strength, they complement each other, together forming a more comprehensive picture of tissue architecture. For instance, changes in gene expression are reflected not only at the molecular level but may also manifest in histological images as morphological alterations (12). Furthermore, the issues of sparsity and dropout in SRT data can be effectively addressed through integrating histological image data (13). By leveraging the interdependence between gene expression and morphological features, as well as the similarity in gene expression patterns among adjacent spots, we can enhance spatial signals and characterize tissue structure. Therefore, achieving a joint representation of multi-modal features in SRT is key to improving data utilization, domain recognition accuracy, as well as functional interpretability.

However, the joint representation of features from different modalities in SRT is challenging. Firstly, these different modalities inherently possess significant heterogeneity. For instance, transcriptomic data are typically high-dimensional, quantified gene expression information, reflecting the gene activity in different spots or cells. In contrast, histology image is two-dimensional visual data, depicting the morphological and structural information of cells at different spots. This fundamental difference makes the direct fusion of these two types of modal difficult. Secondly, the disparity in data scale and resolution is also a crucial issue. Transcriptomic data reveals unique patterns of gene expression within spots or cells from a microscopic perspective. Conversely, histology image provides more macroscopic information on organization and morphology. This difference in scale complicates the establishment of spatial correspondence, thereby posing challenges in comparing and integrating these two categories of data.

Recently, a variety of cutting-edge computational methods have been developed to effectively address the challenge of joint representation of multi-modal SRT data. Specifically, BASS, BayesSpace and Giotto leverage spatial neighborhood information for enhancing the resolution of SRT data (14–16). CellCharter and PRECAST incorporate spatial contexts to correct batch effect for a better domain identification (17, 18). MENDER is a recently proposed multi-range cell context decipherer for ultra-fast tissue structure identification (19). CCST, STAGATE, SpaceFlow and GraphST utilize Graph Neural Networks (GNN) to integrate gene expression data with spatial information, achieving effective clustering of spots (20–23). However, these methods do not employ histology images, failing to fully enhance the interpretability of gene expression data through these images. This limitation often results in subsequent analyses that are less accurate and lack robustness. In contrast, recent pioneering studies like stLearn and DeepST have shown more significant progress (24, 25). These methods effectively integrate gene expression data with spatial neighborhood information and morphological features extracted from histology images, demonstrating a stronger potential for application. Despite these methods demonstrating some capability in processing and interpreting multi-modal information in SRT data, they give less consideration to the complex global spot similarity across high-resolution, content-rich, and distinctive spatial multi-modal features. This limitation impedes their ability to accurately characterize spatial patterns and discover functional biological contexts in tissue.

To achieve precise identification of spatial domains, we introduce stMMR, a geometric deep learning method for effective representing multi-modal information in SRT data. stMMR utilizes spatial location information as a bridge to establish adjacency relationships between spots. It encodes gene expression data and morphological features extracted from histological images using GCN. stMMR proposed a novel strategy to achieve joint learning of intra-modal and inter-modal features. Within a certain modality, stMMR employs self-attention mechanisms to learn the relationships of different spots. For integrating cross-modal information, stMMR innovatively utilizes similarity contrastive learning along with the reconstruction of gene expression features and adjacency information.

To assess the capability of stMMR in representing multi-modal information, we conducted comprehensive tests on various SRT datasets, including samples profiled by 10x Visium, NanoString technology as well as Spatial Transcriptomics (ST) technology. We evaluated the effectiveness of stMMR with several state-of-the-art techniques in domain recognition tasks. The results showed that stMMR achieved significant success in domain identification. Additionally, the analysis of domain-specific genes across multiple datasets revealed that stMMR significantly achieves the enhancement of gene expression data and facilitates the understanding of domain-specific genes. Finally, we conducted thorough analyses on breast cancer datasets based on 10x Visium technology and lung cancer datasets based on NanoString technology (26). Our studies found that stMMR accurately identify tumor edges and tumor-infiltrating regions, demonstrating outstanding continuity in regional identification across different slices, thus proving its potential value in clinical research. Overall, the stMMR method exhibits exceptional capability in the multi-modal feature representation of SRT, providing a powerful new tool for accurate and robust domain identification.

## Materials and methods

### Overview of stMMR

The multi-modal joint representation process of stMMR primarily consists of the following three steps: multi-modal feature embedding, feature fusion and feature reconstruction. The overall workflow of stMMR is illustrated in Figure 1.

**Figure 1.**
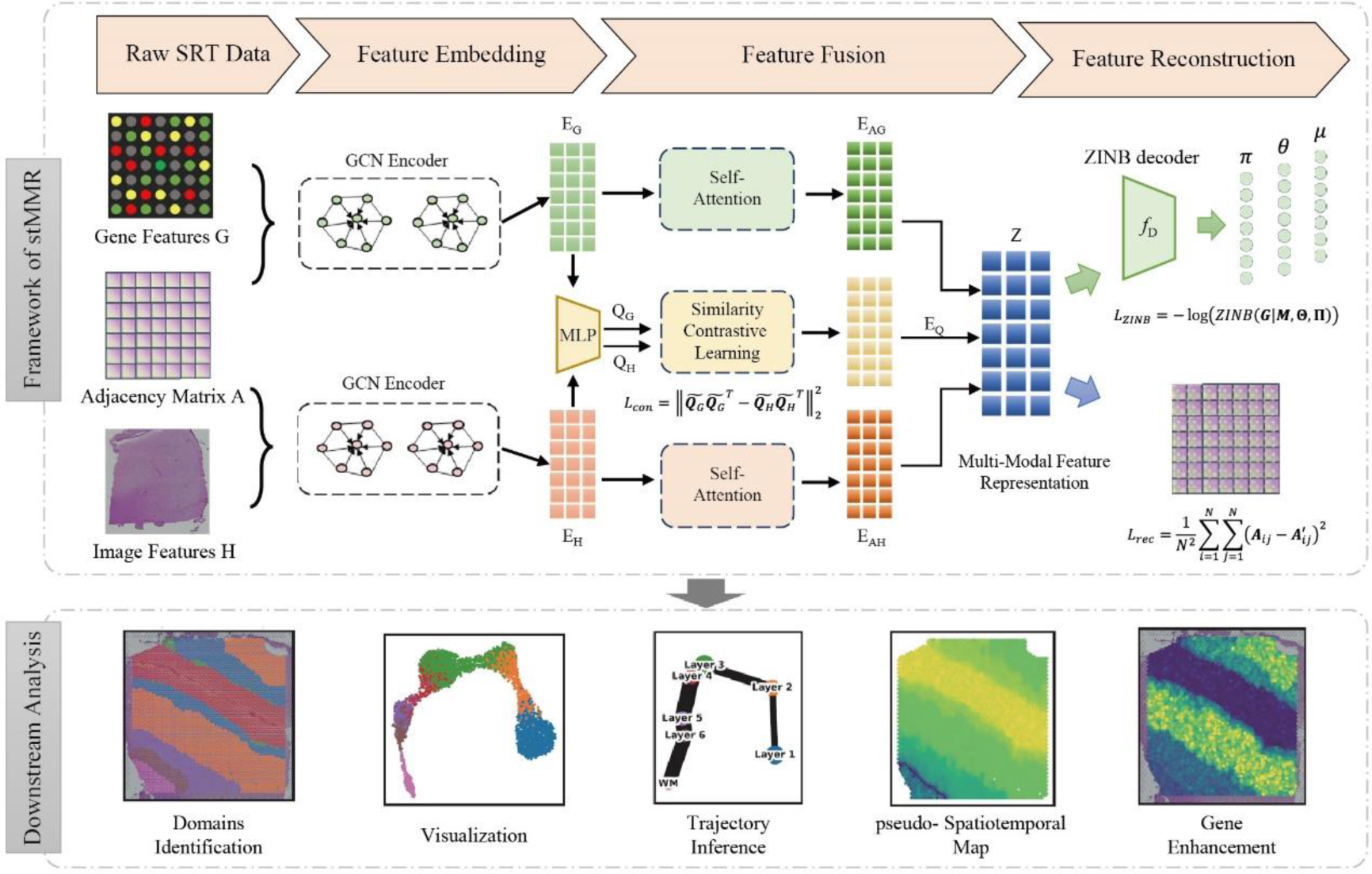
Schematic overview of stMMR for the joint representation of features from different modalities. Gene expression and histology image information are embedded using GCN module based on adjacent matrix. Then, the relationships between different modalities are captured through similarity contrast learning, followed by feature fusion. Finally, the original features are reconstructed from the multi-modal feature representation. This representation can be used for downstream analysis directly.

### Multi-modal feature embedding

The stMMR initially performs embedding on gene expression, spatial location, and histology image information. We begin by assuming the presence of SRT data comprising N spots. For tissue histological images, we extract pixel features corresponding to each spot using a pre-trained Vision Transformer (ViT) model (27), resulting in a feature matrix ***H*** ∈ ℝ^*N*×*M*^, where *M* is the output dimension of the pre-trained ViT model (Supplementary Section 1.3). For gene expression data, we employ SeuratV3 to filter high variance genes and perform a log transformation on the expression levels of these genes, denoted as ***G*** ∈ ℝ^*N*×*P*^, where *P* represents the number of high variance genes identified. Additionally, we encode the spatial location information of each spot, resulting in a position encoding matrix corresponding to each spot. Specifically, we used an undirected weighted graph to present SRT data. For any two spots, we posit that the closer their spatial distance, the greater their similarity. Consequently, we define the adjacency matrix ***A*** between any two spots *i* and *j* as follows:

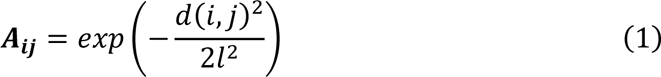

where *d*(*i*, *j*) represents the Euclidean distance between spots *i* and *j*, and *l* is a hyperparameter controlling the relationship between weight and distance. A larger value of *l* implies a faster decay of weight with increasing distance.

GCN is a geometric deep learning module frequently used in graph representation learning in recent years (28). They effectively integrate information from neighboring nodes to achieve efficient representation of target nodes. Subsequently, we employ encoders with two layers of GCNs to perform message passing and aggregation on pixel features and gene expression features, as shown in Eq.2:

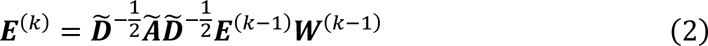

where ***E***^(*k*)^ and ***E***^(*k*−1)^ represent the input and output of the GCN module, respectively. ***E***(^0^) corresponds to the pixel features ***H*** or gene expression features ***G*** for each spot. 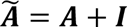 denotes the adjacency matrix of the undirected graph, where ***I*** is the identity matrix. 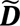 and ***W***^(*k*−1)^ are the weighted degree matrix and trainable parameter respectively. The pixel features and gene expression features obtained after passing through the encoder are denoted as ***E***_*H*_ and ***E***_*G*_.

### Feature fusion

To effectively aggregate multi-modal information, we propose a novel feature fusion strategy. First, stMMR uses a normalized attention module to learn the relationships between spots in a single modality, as shown in Eq.3:

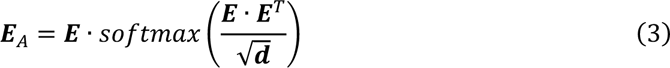

where ***E*** represents the pixel features ***E***_*H*_ or gene expression features ***E***_*G*_ obtained in the previous step. The new features obtained through the attention module are ***E***_*AH*_and ***E***_*AG*_. It is noteworthy that we use a nonlinear activation function and the Euclidean distance matrix ***d*** to normalize the weights. This approach effectively avoids the issue of local optima caused by excessively large weights for certain spots (29).

For cross-modal information, stMMR adopts a contrastive learning approach for feature fusion. Previous research indicates that histology information and gene expression information share both similarities and complementary relationships (30–32). stMMR emphasizes the consistency between multiple modalities by constructing a cross-modal contrastive learning strategy. Specifically, stMMR maps the latent features of multiple modalities, ***E***_*H*_ and ***E***_*G*_, to a space using two fully connected neural networks, thereby obtaining hierarchical representations for both modalities,

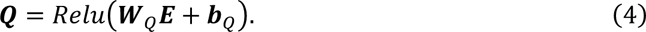

In Eq.4, ***E*** represents the pixel features ***E***_*H*_ or gene expression features ***E***_*G*_, and ***Q*** corresponds to ***Q***_*H*_ or ***Q***_*G*_. ***W***_*Q*_ and ***b***_*Q*_ are the parameters of the fully connected network.

After obtaining the low-dimensional features ***Q***_*H*_ and ***Q***_*G*_ for the two modalities, we further employ a fully connected neural network to fuse these two modalities, as shown in Eq.5:

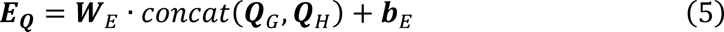

where ***W***_*Q*_ and ***b***_*Q*_ are the parameters of the fully connected network.

To enhance the consistency between ***Q***_*H*_ and ***Q***_*G*_, we use a constraint as shown in Equation 6, replacing the loss of traditional contrastive learning:

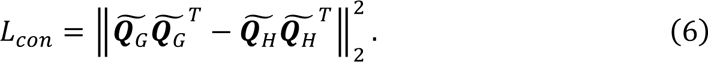

In this equation, 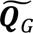 respectively. and 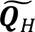 are the normalization matrices of ***Q***_*G*_ and ***Q***_*H*_,

Finally, we further integrate modality specific features ***E***_*AH*_ and ***E***_*AG*_ obtained from Eq.3 with the cross-modality features ***E***_***Q***_ obtained from Eq.5 to get the multi-modal feature representation ***Z***, as shown in the following equation:

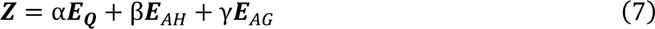

In this equation, α, β, and γ are hyperparameters for adjusting the importance of features.

### Feature reconstruction

Finally, to ensure that the embeddings ***Z*** learned by our model contain biologically information, we designed a reconstruction phase using ***Z*** to reconstruct the original features. Here, we primarily focus on reconstructing the adjacency matrix ***A*** and the gene expression profiles.

SRT data are characterized by high sparsity, discreteness, and variance greater than the mean, specifically manifested as a high number of genes expressed at zero (zero inflation) (33). Previous research has found that the zero-inflated negative binomial (ZINB) distribution can effectively characterize gene expression in SRT (34). Therefore, stMMR also adopts the ZINB to describe gene expression information. In simple terms, we estimate the parameters of the ZINB distribution for each gene (i.e., μ, θ, and π, Supplementary Section 1.4) through three different fully connected networks, as follows:

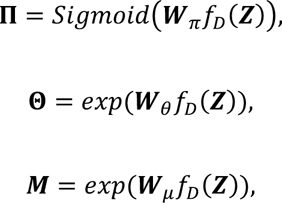

where ***M***, 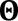, and Π are the matrix forms of μ, θ, and π, representing the mean, dispersion, and dropout probability of the output from network respectively. The dropout probability ranges between 0 and 1. Due to the non-negative nature of the mean ***M*** and dispersion 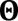, we use the exponential function, and for the dropout probability Π, the Sigmoid function is used. *f*_*D*_ is a decoder with a fully connected layer.

Based on the aforementioned reconstruction of gene expression information, we further designed the loss function for the ZINB decoder, as shown below:

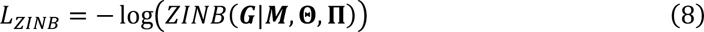

For spatial neighborhood relationships, we adopted the concept of a graph auto-encoder to directly estimate the adjacency matrix (35, 36), as shown in Eq.9:

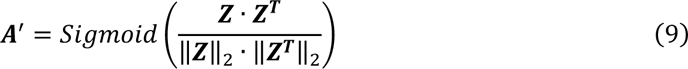

Subsequently, we defined a function to calculate the regularization loss between the reconstructed matrix and the adjacency matrix, as in Eq.10:

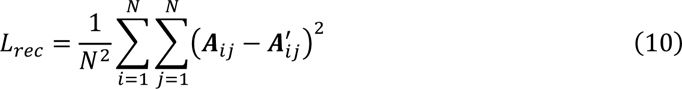

This function quantifies the difference between the original adjacency matrix ***A*** and its reconstructed version ***A***^′^, thereby facilitating the accurate reconstruction of spatial relationships.

### Objective function

Finally, we integrated Eq.6, Eq.8, and Eq.10 to formulate the final objective function, as shown in Eq.11:

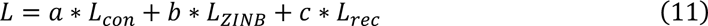

In this equation, *a*, *b*, and *c* are hyperparameters that control the contribution of the different loss terms.

For detailed information on the training process and parameter settings, please refer to the Supplementary Section 1.5 and 1.6.

### Datasets

All the dataset used in this study are listed in Supplementary Table S1. For detailed descriptions and processing procedures, please refer to Supplementary Section 2. Processed datasets are also available at SODB and can be loaded by PySODB (37, 38).

### Benchmark methods

To demonstrate the effectiveness of the multi-modal feature representation in SRT data, we selected 7 different state-of-the-art methods for benchmarking comparison.

These methods include SCANPY (39), which utilizes only gene expression data; CCST, STAGATE, GraphST, and SpaceFlow, which employ both gene expression and spatial location information (20–23); and stLearn and DeepST, which incorporate all three modalities (24, 25). Methods that have already been compared in previous works are not included in our analysis (40–42).

## Results

### stMMR enhances detection of stratified architectural patterns in human dorsolateral prefrontal cortex (DLPFC) tissue

The spatial structure of the brain is closely related to its function, particularly evident in the layered organization of the human brain cortex. Cells in different cortical layers not only exhibit unique gene expression patterns but also vary in morphology, physiology, and connectivity (43). To explore the spatial structure arrangement of brain, we collected a 10x Visium dataset containing 12 dorsolateral prefrontal cortex (DLPFC) sections (44). Maynard et al. manually annotated different layers (layer 1-6) and white matter (WM) of the DLPFC based on morphological features and gene markers (Figure 2A) (44). This annotation served as the ground truth for comparing and analyzing the effectiveness of stMMR and other advanced spatial domain identification methods.

**Figure 2.**
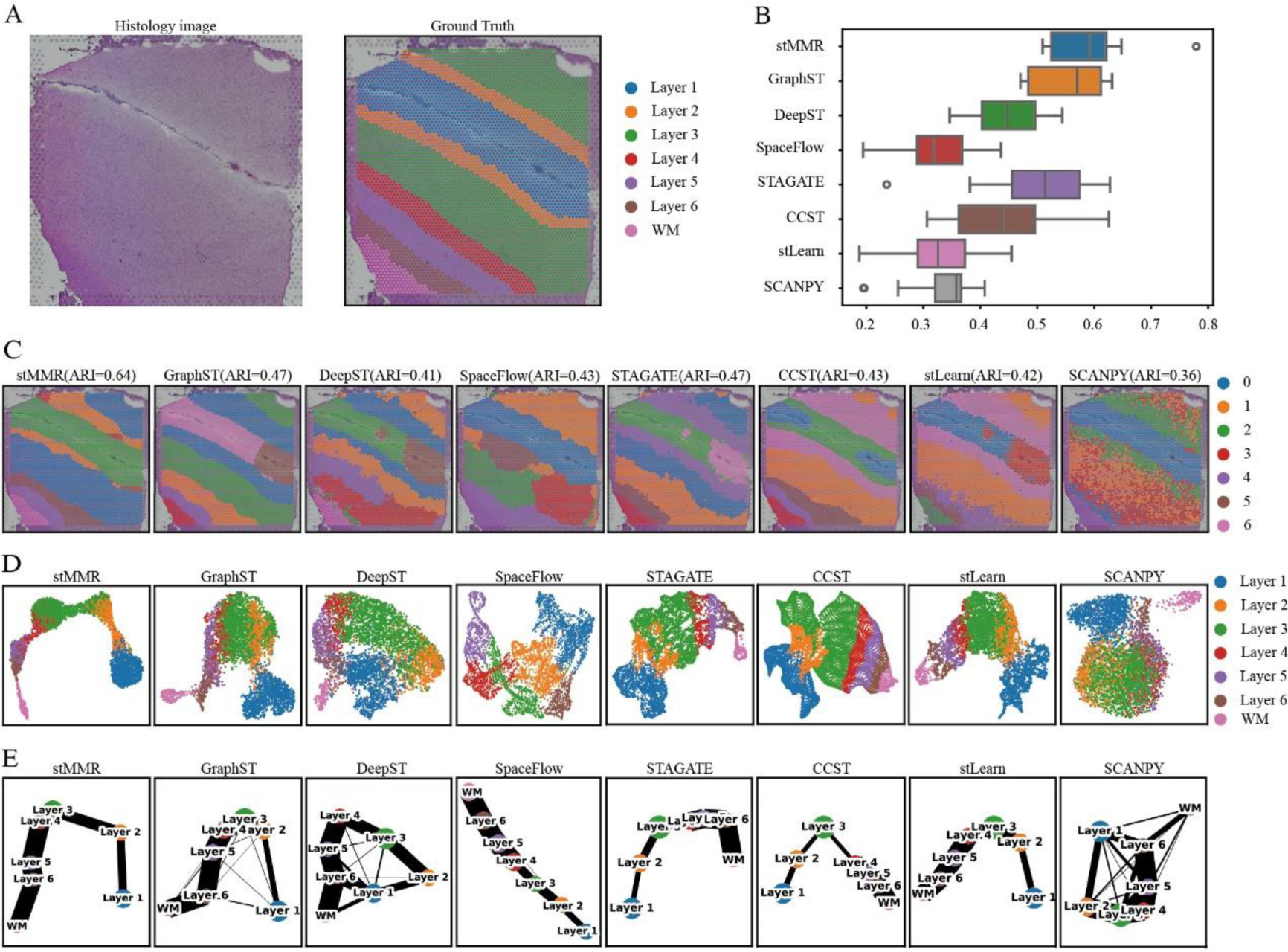
Performance comparisons of different methods on DLPFC datasets. (A) The histology image and manually annotated brain regions of slice 151509. (B) The overall performance of 8 different methods across 12 slices. (C) The domain recognition results on slice 151509. (D) The UMAP visualization results of the embeddings from 8 different methods on slice 151509. (E) The inferenced trajectories on slice 151509.

We initially compared the Adjusted Rand Index (ARI) levels of various methods across 12 slices of the DLPFC dataset, as shown in Figure 2B. The results reveal that stMMR outperformed other methods, achieving the highest ARI compared to manual annotations. This result also demonstrates that stMMR possesses the smallest variance across all slices. Notably, the results from STAGATE, CCST, and SpaceFlow show differences in the ARI across different slices, indicating that these methods are more sensitive to the difference of domains. Scanpy uses only gene expression information and shows the poorest performance. Methods like stlearn and DeepST, which integrate histological information, are outperformed by GraphST and STAGATE. This underperformance might stem from insufficient integration of transcriptomic and imaging data.

Next, we conducted a detailed analysis for each slice (Figure 2C and Supplementary Figure S1). To demonstrate the results, we used slice 151509 as an example (Figure 2C-E). The results shows that DeepST struggle with rough segmentation between layers. CCST, SpaceFlow, and stLearn have issues with erroneous region identification. Although GraphST and STAGATE accurately discern the arrangement of different regions, these methods exhibit biases in identifying the boundaries between distinct domains. In this specific case, stMMR demonstrates exceptional domain identification results. We further utilize UMAP for low-dimensional visualization analysis of the results obtained from different methods (Figure 2D), to verify whether the embeddings can accurately encompass information on regional arrangement and boundaries. The analysis reveals that techniques such as stMMR, CCST, STAGATE, and stLearn effectively separate different domains. In contrast, GraphST, SpaceFlow and DeepST exhibit noticeable issues in layer boundaries. For instance, the boundaries between layers 2, 3, and 4 are confused.

Further, we conducted a detailed trajectory inference using the PAGA algorithm (45) for these methods (Figure 2E). The PAGA graphs indicates that stMMR, STAGATE, CCST, and stLearn performs well in predicting trajectory between adjacent layers. The other methods display confused results in this analysis.

Moreover, we identified domain-specific genes in the human brain cortex using differential expressed gene analysis. Genes such as AQP4 and HPCAL1 are recognized as layer-specific genes (Supplementary Table S2). These genes are enriched in multiple layers and have been confirmed through multiplex single-molecule fluorescent in situ hybridization (44).

Combining the insights from these various analyses, it is evident that stMMR remarkably effective in several critical tasks, including domain identification, pseudo-spatiotemporal analysis and domain-specific genes discovery. These results adequately demonstrate effective capability of stMMR in integrating transcriptomic and histological data.

### stMMR enhances spatial gene expression profiling and structural characterization

In SRT, the analysis of domain-specific genes holds significant importance. The functionality of complex tissues is closely related to distribution domain-specific genes. However, in SRT research, identifying domain-specific genes which have relationships with histological structures is challenging. This is primarily due to the presence of substantial noise in the gene expression profiles generated by SRT techniques. The noise mainly arises from technical issues during sequencing, such as the dropout event (46–48). To validate whether stMMR can enhance gene expression data through histological information, we analyzed domain-specific gene expression in 151509 slices of the DLPFC dataset. Inspired by the previous study (49), we validated the effectiveness of the stMMR in enhancing gene expression data by comparing the relationship between the original gene expression profile and the profile reconstructed through the ZINB decoder with manually annotated regions.

Compared to using original gene expression data, employing reconstructed gene expression data facilitates the identification of a larger number of domain-specific genes across all brain layers (Supplementary Table S2). For instance, when using the original gene expression data, the CACNA2D2 and ADCYAP1 were not identified as domain-specific genes in layer 3. However, after enhancement with stMMR, we were able to accurately detect the CACNA2D2 and ADCYAP1 as domain-specific genes in layer 3. Notably, previous research has found that in layer 3 of primates, the CACNA2D2 gene exhibits differential expression and is closely associated with several biological pathways, including calcium signaling and synaptic long-term depression (50). ADCYAP1 has also been proved to be a domain-specific gene in former study (51). This suggests that the expression patterns after stMMR enhancement are more consistent with known neurobiological functions and pathological states.

We also conducted a more detailed analysis by combining gene expression levels with their spatial locations (Figure 3). We found that after enhancing gene expression with stMMR, more distinct expression patterns of domain-specific genes can be observed between brain layers. Specifically, Figure 3A demonstrates a clear spatial representation of domain-specific marker genes (ADAYAP1, CACNA2D2, CALB1, MARC1, MB and LPL) after data enhancement. For example, in the original data, the expression pattern of genes such as ADCYAP1, CACNA2D2 and MB are sparse, and the boundaries in spatial regions are blurred, making it difficult to discern a clear expression pattern (Figure 3A and B). However, after enhancement with stMMR, we can observe that CACNA2D2 and CALB1 exhibit much clearer expression patterns in layers 3 and 4. Additionally, the enrichment of MARC1, MB, and LPL in the white matter regions become more pronounced (Figure 3A and C). These results reflect that stMMR not only improves the spatial resolution of gene expression patterns using histological information but also enhances our understanding of the subtle differences in gene expression across different regions of the brain.

**Figure 3.**
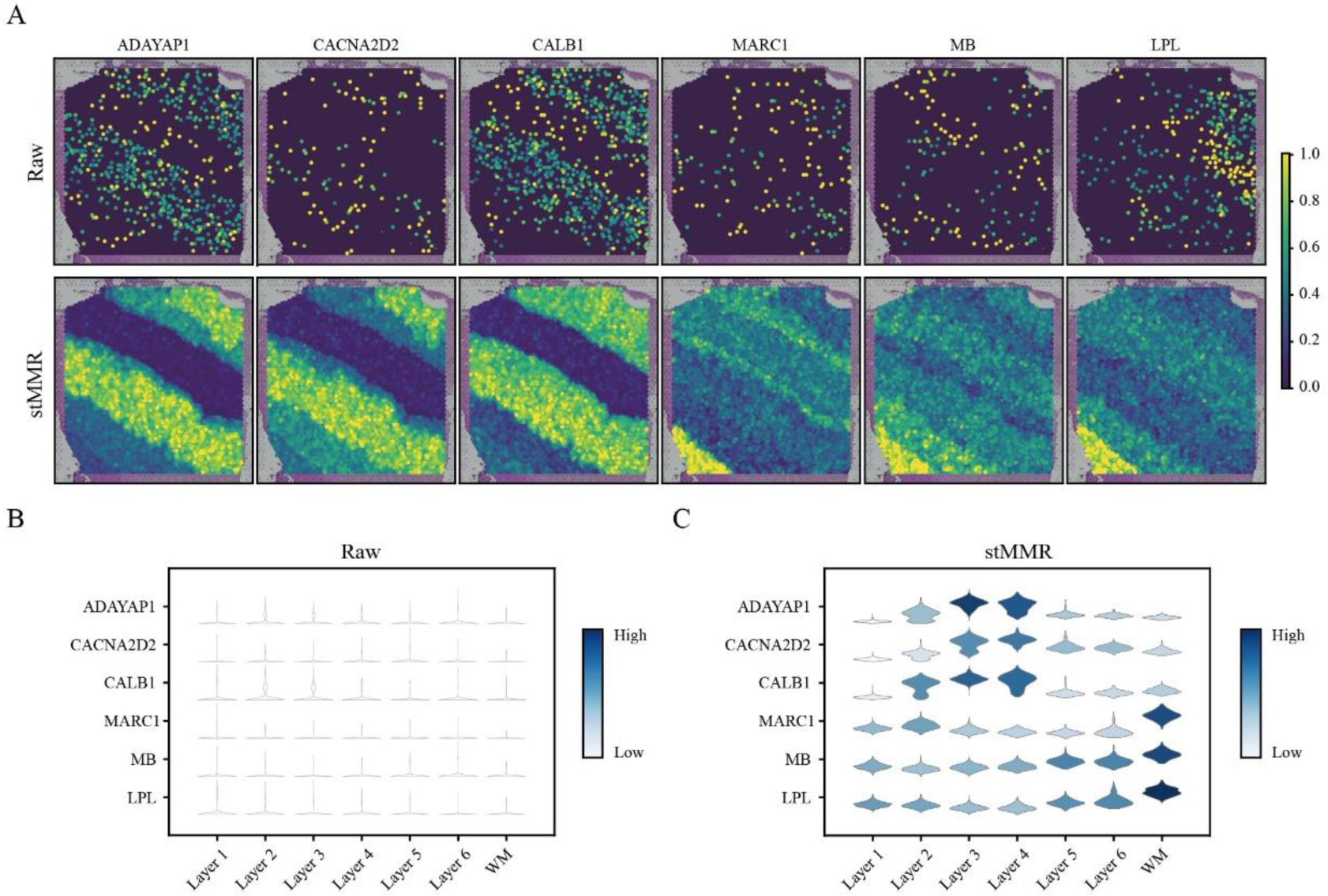
stMMR enhances spatial gene expression profiles and spatial structural characterization. (A) Spatial representation of layer-specific marker genes before and after data enhancement. (B) Gene expression level before and after data enhancement.

### stMMR deciphers evolving cell lineage structures in chicken heart ST dataset

Analyzing temporal SRT data can reveal the dynamic domain changes during the development of tissue organs. This is crucial for our understanding of complex biological processes such as tissue development and disease progression. We collected the chicken heart SRT dataset to further investigate the effectiveness of stMMR in the integrated representation of multi-modal features (52). This dataset includes 12 tissue slices, collected on day 4 (5 slices), day 7 (4 slices), day 10 (2 slices), and day 14 (1 slices), documenting four key stages of the Hamburger-Hamilton ventricular developmental stages (52).

We annotated the slices of different developmental stages using labels provided by the original research (Figure 4A) (52). Subsequently, we employed the embeddings from stMMR and SpaceFlow to identify domains of chicken heart across these four distinct stages. Figure 4B indicate that the regions detected by stMMR largely coincide with manual annotation. For instance, in all tissue sections, major regions of the chicken heart, such as atrial cells and the inter-ventricular septum, are accurately identified. This discovery is significant for a deeper understanding of the spatial structure of cardiac tissues. Notably, stMMR also detects domains that are hard to identified (Figure 4B). For example, in the data from days 7, 10, and 14, the epicardium, a thin layer surrounding the outer side of the chicken heart, is clearly identified by stMMR. Although there are some instances of misclassification in the characterization of spot features using stMMR in a few regions, the identification of the epicardium is quite clear (Figure 4B).

**Figure 4.**
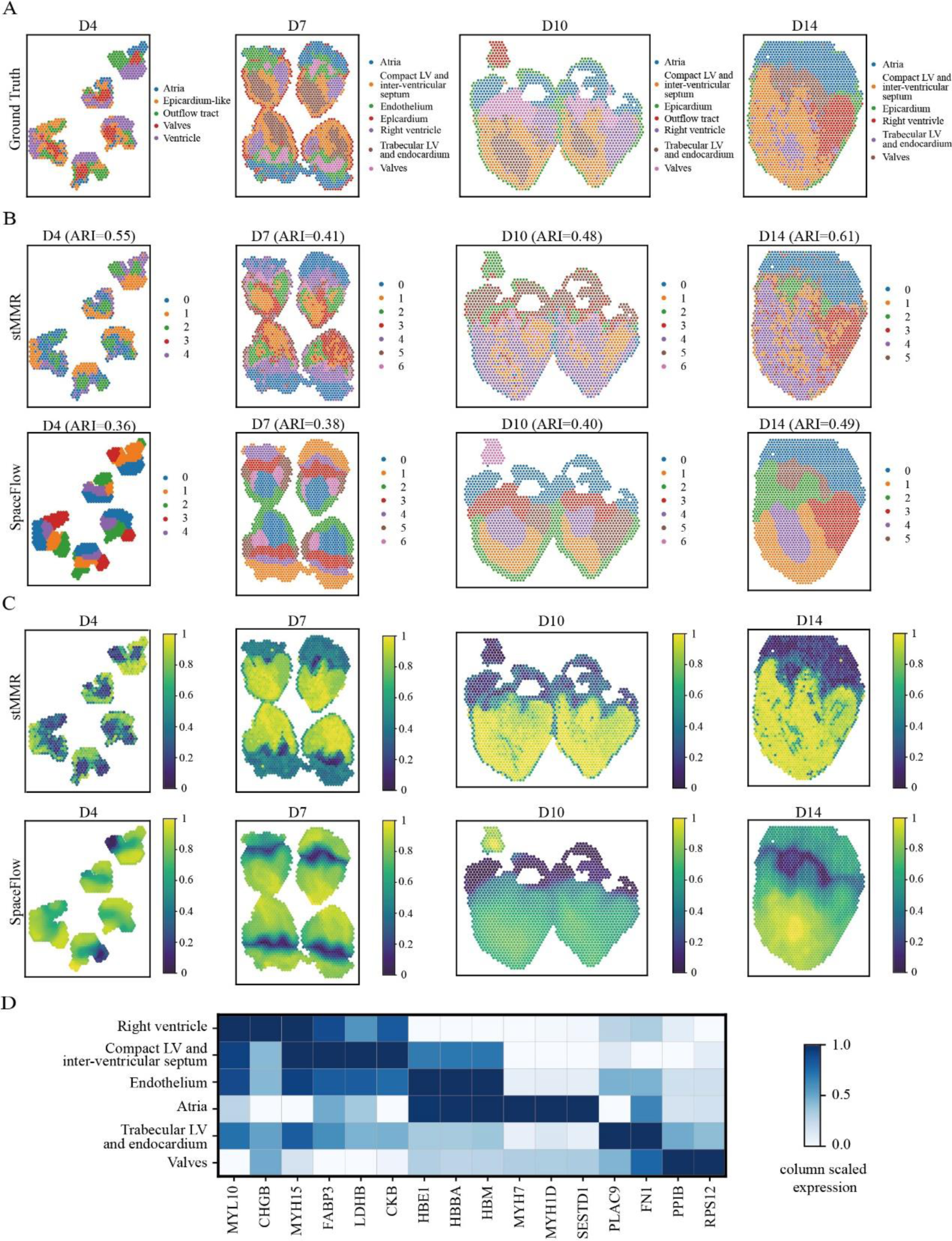
stMMR reveals cell lineage structures during chicken heart development. (A) The ground truth label provided by the original data. (B) The domains recognized by stMMR and SpaceFlow. (C) The plots of pSM value from stMMR and SpaceFlow for illustrating pseudo-temporal developmental trajectory. (D) The differentially expressed marker genes discovered by stMMR.

Next, we adopted a method similar to previous study to analyze the pseudo-spatiotemporal map (pSM) (23). In brief, we mapped the spot features obtained through stMMR and SpaceFlow on the pseudo-temporal axis (23, 39, 53). These points reflect the relative positions of cells in their developmental trajectory or functional state. As clearly visible in Figure 4C, within the D7 to D14, the valve structures can be distinctly identified through the pSM values. Moreover, the representation of the myocardium in ventricles, as indicated by the pSM values, appears more uniform (yellow area) compared to the regional segmentation results in Figure B. According to related research (54), the endocardium, the inner layer of the heart, is one of the early events in cardiac formation. The endocardial tubes are fundamental to cardiac development, eventually merging to form the primitive heart tube. As the heart tube forms, myocardial development commences, followed closely by the development of the atria. In our analysis, we observe that the myocardium in ventricles (yellow area in Figure 4C) consistently shows higher pSM values compared to other areas in the same stage, indicating a later pseudo-temporal ordering of the ventricular myocardium (23). Additionally, the pseudo-temporal ordering of the atria (marked in teal) follows that of the valves, suggesting that the development of the atria occurs after the valves. Therefore, the pSM derived from stMMR accurately displays the developmental sequence of the chicken heart. We further identified domain-specific genes through differential expression analysis across regions. For instance, we observed that MYH7 is highly specifically expressed in the Atria. This finding aligns with previous reports on the analysis of Atria and Ventricles specific proteins (55).

In addition, we also employed various algorithms to perform a comparative analysis of the pSM on the human DLPFC dataset. As shown in Supplementary Figure S2, we observe that in the pSM analysis, methods like GraphST, DeepST, stLearn, and SCANPY fail to clearly reflect the layered spatial organization of the tissue. In contrast, the stMMR, STAGATE, SpaceFlow and CCST are able to distinctly reveal the layered pattern of the pSM, displaying clear and smooth color gradients. This result from stMMR not only mirrors the correct internal and external developmental sequence of the cortical layers but also demonstrates the layered spatial organization of tissue, aligning with the findings of previous studies (23).

### stMMR accurate identifies tumor region in human breast cancer

Breast cancer is a major type of cancer worldwide (56). We collected human breast cancer SRT data from the 10x Visium platform to conduct an in-depth study of the microenvironment in breast cancer. This dataset includes 3, 798 spatial spots and 36, 601 genes. Experienced pathologists have annotated these SRT data using H&E images and signature genes of breast cancer, categorizing them into 20 distinct regions (Figure 5A).

**Figure 5.**
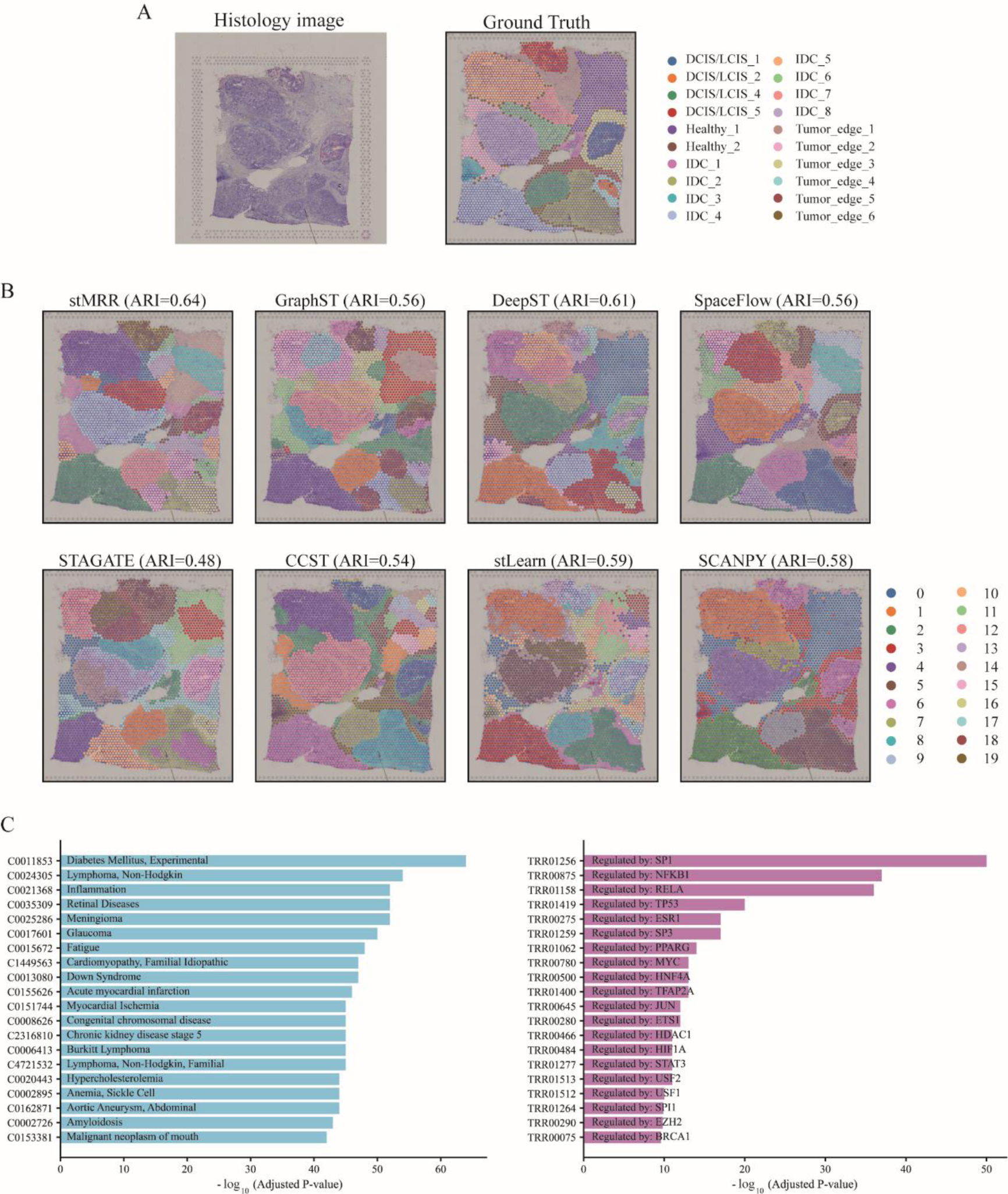
stMMR identifies tumor region in human breast cancer dataset. (A) The H&E images and the manually annotated regions. (B) The annotation results from different methods. (C) Top 20 differentially expressed gene enriched terms identified by DisGeNET (left panel) and TRRUST (right panel).

First, we applied different methods for domain identification. From the results presented in Figure 5B, it is observed that stMMR shows the most outstanding performance in category labeling. In terms of regional continuity, stMMR also demonstrates superior performance among different methods. Taking the IDC_5 area in the upper left corner as an example, this area occupies a significant portion in invasive ductal carcinoma, with a notable increase in cancer cells compared to normal tissue or non-tumorous areas. Previous studies have also indicated that cells originating from solid tumors are primarily concentrated in the IDC area (22). However, only stMMR accurately identified the entire IDC_5 area, demonstrating higher precision compared to other methods. Additionally, stMMR also exhibits higher continuity in predicting the Tumor_edge area, whereas the results of other methods appear more dispersed in this aspect.

Next, we conducted a comprehensive analysis of domain-specific genes between merged tumor and normal regions (Supplementary Section 2.3). We utilized the DisGeNET to delve into the differentially expressed genes in the breast cancer tumor regions (57). Our analysis revealed that these domain-specific genes are enriched in several breast cancer related terms (Figure 5C left panel). For instance, C0024305 is a term related to non-Hodgkin lymphoma. Studies have shown that the development of breast cancer significantly increases the risk of non-Hodgkin lymphoma, particularly follicular lymphoma and mature T/NK cell lymphomas (58). This risk is notably more pronounced in patients undergoing hormone therapy and in younger patients (58). Importantly, genes were also enriched in an inflammation-related term (Figure 5C left panel). Numerous studies have indicated that inflammation plays a regulatory role in the development of cancer and its response to treatment (59–61). To further validate our research findings, we conducted a systematic analysis of the transcriptional regulatory network using TRRUST (62). The analysis results indicated that multiple top-ranked terms are closely associated with breast cancer (Figure 5C right panel). For instance, TRR01419 emerges as the fourth most significant term, with TP53 is identified as its key regulatory factor. TP53 plays a crucial role in both cancer-related systemic inflammation and the progression of cancer (63). Additionally, the key regulatory factors for the top three terms - TRR01256 (regulated by SP1), TRR00875 (regulated by NFKB1), and TRR01158 (regulated by RELA) - have also been confirmed in previous studies to play pivotal roles in the development and progression of breast cancer (64–68).

### stMMR dissects cell type differences in a lung cancer SRT dataset based on NanoString technology

To further validate the generalization ability and applicability of stMMR, we tested its effectiveness using the single-cell SRT dataset generated by NanoString CosMx SMI. This dataset comprises lung cancer tissue samples from 20 fields of view (FOVs) (26), involving 982 genes and 83, 621 cells, covering eight major cell types, including lymphocytes, neutrophils, mast cells, endothelial cells, fibroblasts, epithelial cells, myeloid cells, and tumor cells. For ease of observation, we displayed the ground truth of one of the FOVs in Figures 6A.

**Figure 6.**
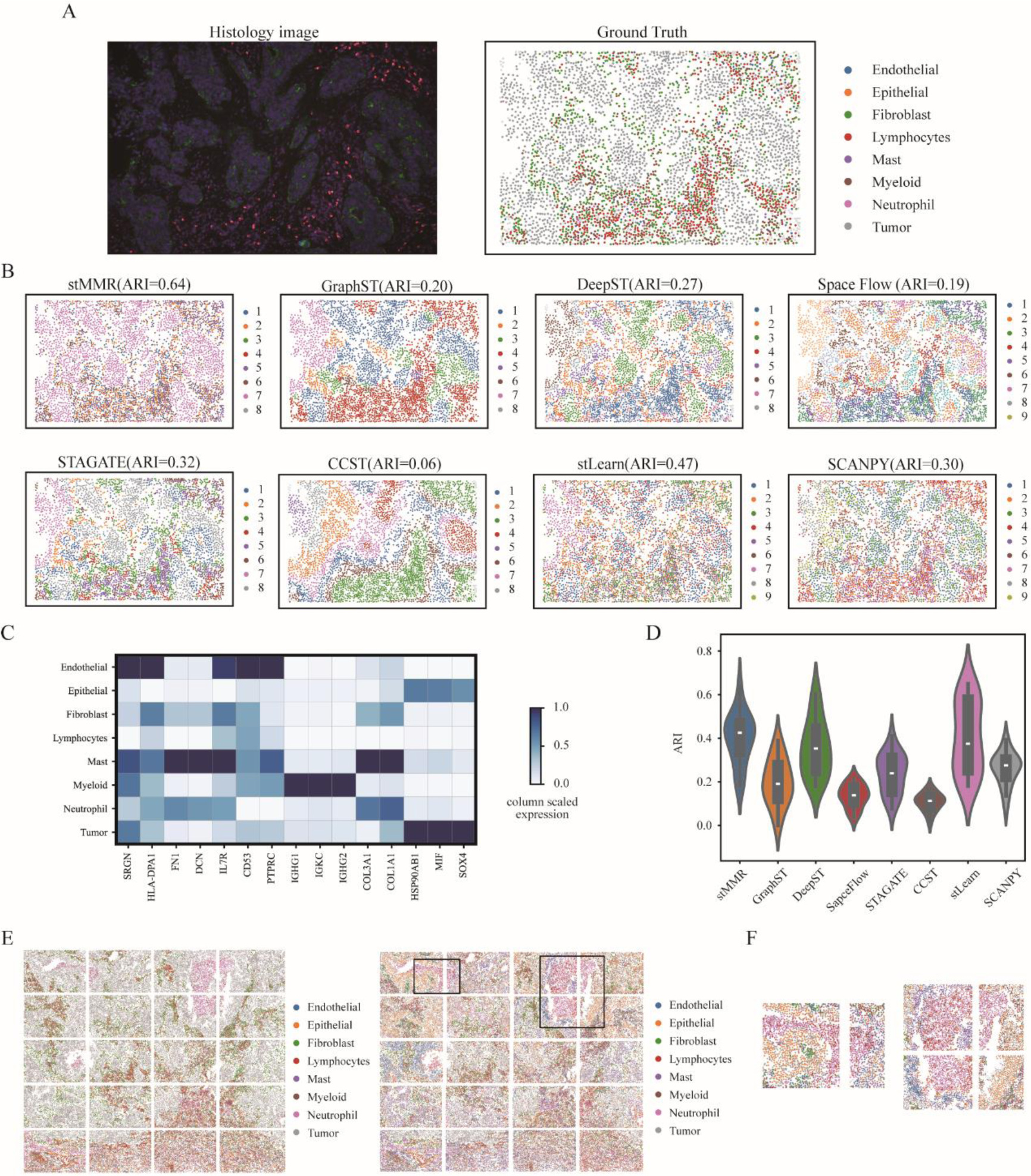
stMMR recognizes cell type differences in lung cancer dataset. (A) One FOV of the lung cancer SRT data. (B) Cell types identified by different methods. (C) Expression pattern of marker genes for different cell types. (D) The overall performance of different methods across 20 FOVs. (E) Cell types annotated manually in 20 FOVs. (F) Cell types annotated by stMMR in 20 FOVs. (G) The zoomed-in results of boundaries between adjacent FOVs identified by stMMR.

We employed the benchmarking methods to conduct a detailed analysis of the spatial organization within 20 FOVs, as shown in Figures 6B, D-F. Figure 6B revealed that stMMR closely aligns with the original study in detecting the spatial distribution of cell types. Particularly in identifying tumor cells, stMMR demonstrates high precision, accurately detecting tumor cells distributed across different locations in the tissue sections (Figure 6B). In the overall analysis of the 20 sections, the performance of stMMR is superior to other methods (Figure 6D). Furthermore, we conducted a cell type-specific gene analysis based on the cell annotations in one slice. We observe that different tissue cells exhibit unique expression patterns (Figure 6C). For instance, Igkc transcripts, previously reported to be upregulated in myeloid progenitor populations, is also confirmed in our study (69). The genes COL3A1 and COL1A1 shows significant positive correlations with neutrophils (70, 71). Additionally, the oncogene SOX4 is prominently featured in our differential analysis of tumor cells (72). These genes are also identified as diagnostic or prognostic biomarkers in previous studies (70, 73–76). Notably, some cell types also share similar gene expression patterns (Figure 6C). For example, epithelial cells and tumor cells in this lung cancer dataset exhibit expression similarities. Multiple studies using single-cell transcriptomics analysis have revealed that lung cancer cells share characteristics similar to those of Type 1 (AT1) and Type 2 (AT2) alveolar epithelial cells (77, 78). This similarity may be related to lung cancer cells maintaining epithelial cell functions, such as cell adhesion and migration (79, 80).

We also conducted a visualization analysis comparing the results of stMMR applied to 20 tissue sections with the actual division of tissue regions. The analysis demonstrates that stMMR effectively identifies tissue regions across multiple sections (Figure 6E). Notably, even in regions bisected by section boundaries, stMMR maintains smooth and continuous (Figure 6E and F). These findings indicate that the joint representation of stMMR not only effectively eliminates noise from different data types but also maintains excellent performance in the recognition of tissue regions across multiple slices.

## Discussion

SRT technology enables us to deeply understand the spatial structure of tissues within biological systems from multiple dimensions, including gene expression profiles, spatial positioning, and histological information. Through comprehensive integration of these modalities, we can obtain an informative joint representation. However, the inherent data heterogeneity along with the varying spatial resolutions presents challenges in the integration of these modalities. To overcome this problem, we propose a novel computational framework, stMMR. This framework aims to harmonize and unify multi-modal data as well as achieve effective joint representation for multi-modal SRT data.

stMMR effectively unifies gene expression profiles and histological information by utilizing spatial location as a connecting link. This method automates the construction of adjacency relationships between neighboring spots. Then, GCN is employed to extract features from both gene expression profiles and histological information. Furthermore, stMMR adopts an innovative strategy for representing intra-modal and inter-modal features. Initially, it employs an attention mechanism for an in-depth learning within a single modality. It then integrates cross-modality features through a combination of similarity contrastive learning, along with the reconstruction of gene expression and adjacency relationship. By applying stMMR to SRT data of various tissues and resolutions, we have validated its exceptional performance in multiple analyses, including domain identification, pseudo-spatiotemporal analysis, gene expression data enhancement as well as the identification of domain-specific genes.

The remarkable performance of stMMR can be attributed to several innovative designs. The most crucial aspect is the integration of histological information with gene expression data through spatial location. In SRT, gene expression data suffers from issues of sparsity and zero inflation, which are key factors that interfere with downstream analysis (33, 81). Previous research has shown that histological information can predict gene expression data (30–32). Therefore, compared to methods that rely on gene expression information solely, stMMR integrates imaging information and exhibits superior performance in spot characterization. Secondly, unlike other methods that construct spatial transcriptomic data as undirected, unweighted graphs, stMMR builds undirected weighted graphs inversely proportional to Euclidean distances between spots, better reflecting the influence of spatial distance on message passing and aggregation. Furthermore, the consideration of relationships within and between modalities is also crucial. Sole reliance on gene expression data for correlation analysis may result in information loss. In contrast, methods that incorporate imaging information, such as DeepST, focus primarily on the integration of multi-modal data, overlooking the relationships within individual modalities. To fully leverage the relationships within and between modalities, stMMR not only uses similarity contrastive learning for integrating features across modalities but also incorporates a self-attention module for deep embedding of features within a modality. Additionally, the reconstruction modules for gene expression and adjacency matrix further encourage the model to retain as much original information as possible. This encoder-decoder structure improves the ability of stMMR to recover information also endows stMMR with denoising capabilities and robustness.

It is noteworthy that the stMMR framework, distinguished by its exceptional feature embedding capabilities, also exhibits a remarkable ability to handle data derived from diverse experimental techniques. Beyond the previously mentioned datasets, our investigation also extended to the analysis of a mouse brain dataset derived using 10x Visium technology and a human pancreatic ductal adenocarcinoma dataset obtained through ST technology (Supplementary Figure S3 and S4). In these tests, the stMMR consistently achieved optimal results.

stMMR also has model scalability. In the design of the model, we considered that gene expression of each spot corresponds to a small area in space. Therefore, before embedding histological information, we also tailor it to match each spot aera. This means that the input information of stMMR is always associated with each spot area. Recently, the advancement of spatial multi-omics technologies has provided new data for analyzing the spatial distribution and functions of cells in tissues from multiple perspectives, such as simultaneous observations of transcriptomes and proteomes, transcriptomes and epigenomics (82–85). The stMMR framework can be easily expanded to support these types of data. The researchers only need to duplicate the gene expression module and then apply similar methods for embedding features within and between modalities. Moreover, the joint representation obtained through stMMR can also be applied to other tasks, such as cell type deconvolution (86–89). This application requires a process similar to methods like scaden (90), where a neural network is connected to the joint representation. Subsequently, the generation of simulation data and model training can be conducted using annotated single-cell data.

There is still room for the improvement of stMMR. Currently, stMMR employs Euclidean distance in the construction of spot adjacency matrices. However, in practical scenarios, it may be more rational to utilize different distance metrics for graph construction based on modal features. For instance, considering gene expression data, the use of Pearson Correlation Coefficients or K-L divergence might be more appropriate to measure expression similarity between spots. In contrast, for spatial imaging data, either Euclidean distance or staining similarity can serve as the distance metric. Under these circumstances, the constructed graph transitions from being a homogenous graph to a heterogeneous one. For such heterogeneous graphs with multiple types of edges, we can apply methods like metapath2vec or multi-view learning to achieve embedding and integration of different modalities (91–94).

In this paper, we introduce a robust and accurate tool, stMMR, for the integration of gene expression data, spatial information, and histological information from SRT data. Compared to existing methods, stMMR demonstrates a significant advantage in integrating multi-modal data, particularly excelling in domain identification, pseudo-spatiotemporal analysis, and domain-specific gene analysis. Overall, as an effective and user-friendly tool, stMMR enhances the multi-modal joint analysis of SRT data, providing substantial support for research in relevant fields.

## Data availability

All datasets used in this paper are publicly available. The descriptions and download address are listed in Supplementary Section 2.1 and Supplementary Table S1. Processed datasets are also available at SODB (https://gene.ai.tencent.com/SpatialOmics/) and can be loaded by PySODB (https://protocols-pysodb.readthedocs.io/en/latest/). The source code for stMMR can be downloaded from Github (https://github.com/nayu0419/stMMR).

## Supplementary data

Supplementary Data are available at NAR online.

## Author contributions

R.G., W.Z. and Z.Y. conceived and supervised the project. D.Z. and N.Y. designed the model and developed the stMMR software. D.Z., N.Y. and W.Z. wrote the manuscript. X.S. and Q.Z. collected and constructed the benchmark datasets. W.L. X.L. and Z.L. conducted biological interpretation. R.G., W.Z. and Z.Y. reviewed the manuscript. All authors read and approved the final manuscript.

